## Supplementary figures and images for "Ribosome subunit attrition and activation of the p53–MDM4 axis dominate the response of MLL-rearranged cancer cells to WDR5 WIN site inhibition"

### Figure 3 Source Data 3

**Figure 3- source data 3**

MB 680R

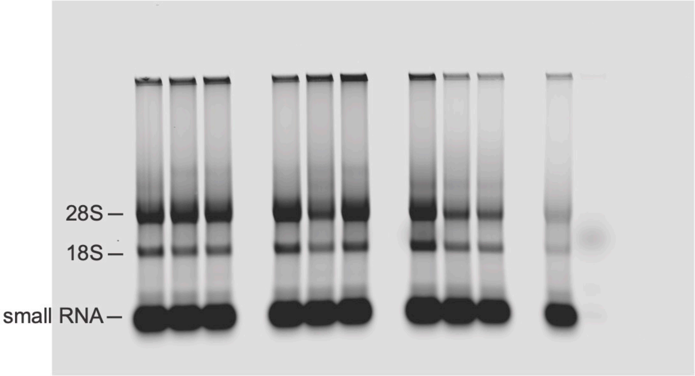


SYBR Safe

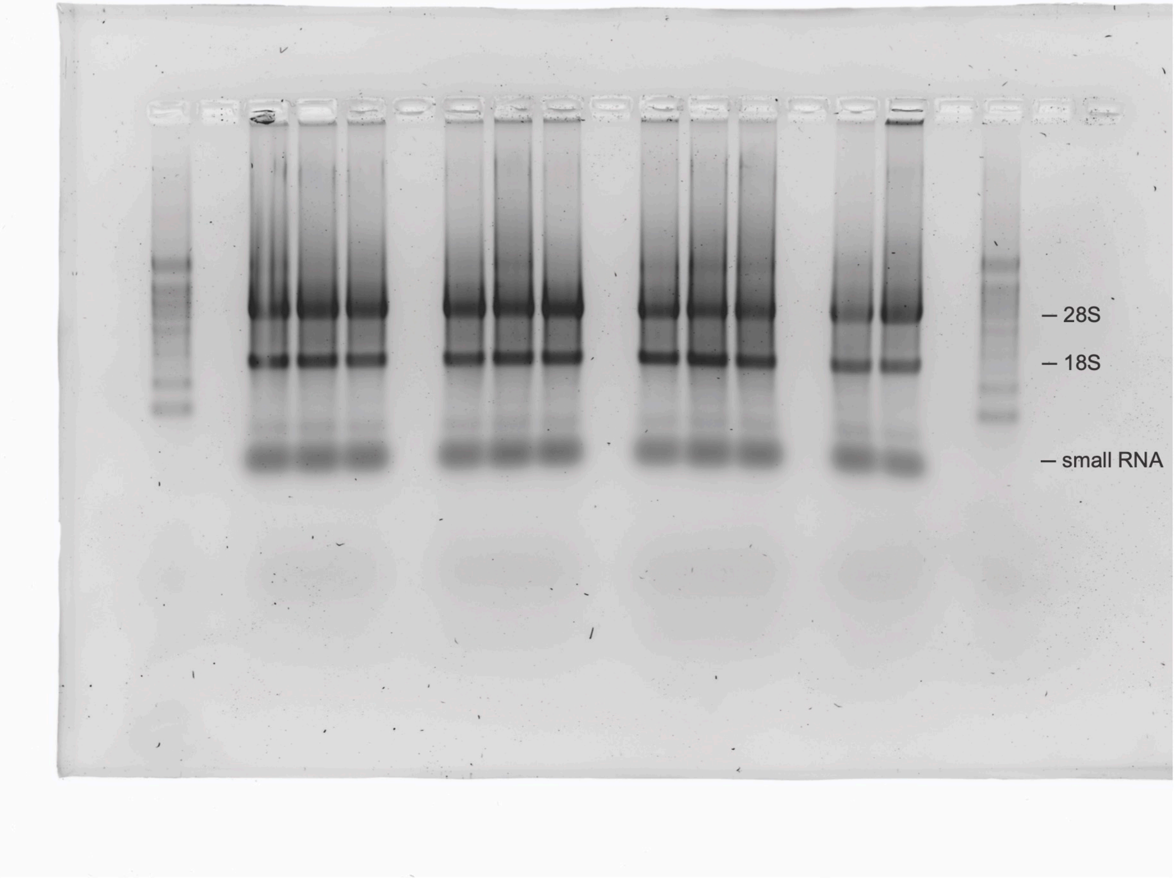

### Figure 6 Source Data 1

#### Figure 6—source data 1

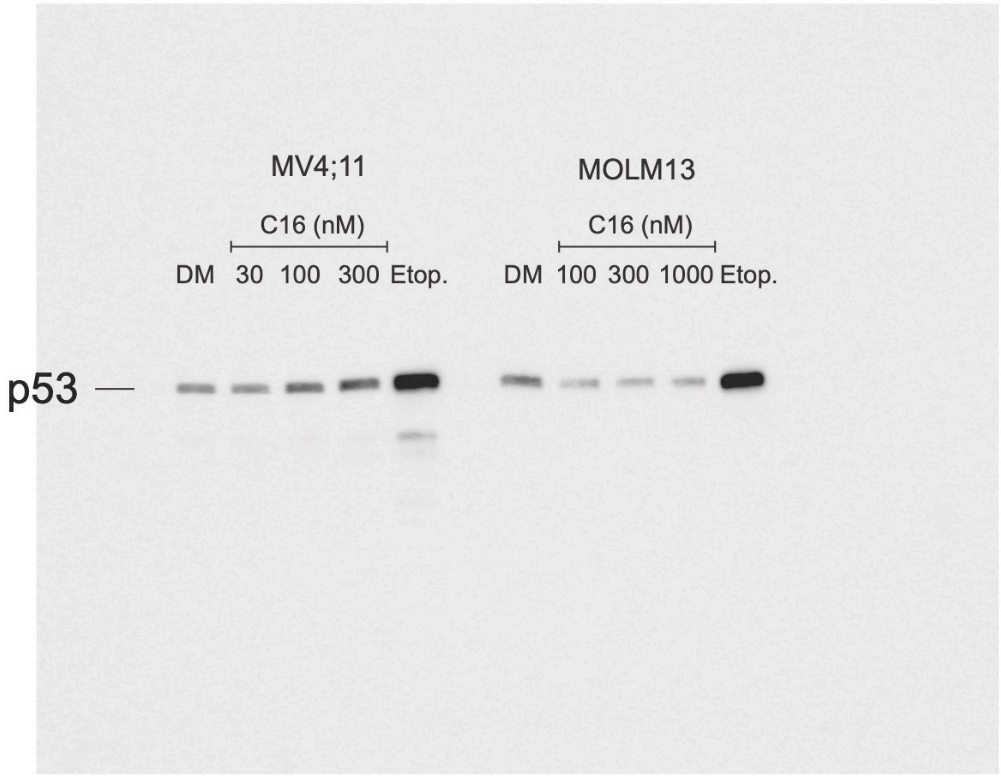


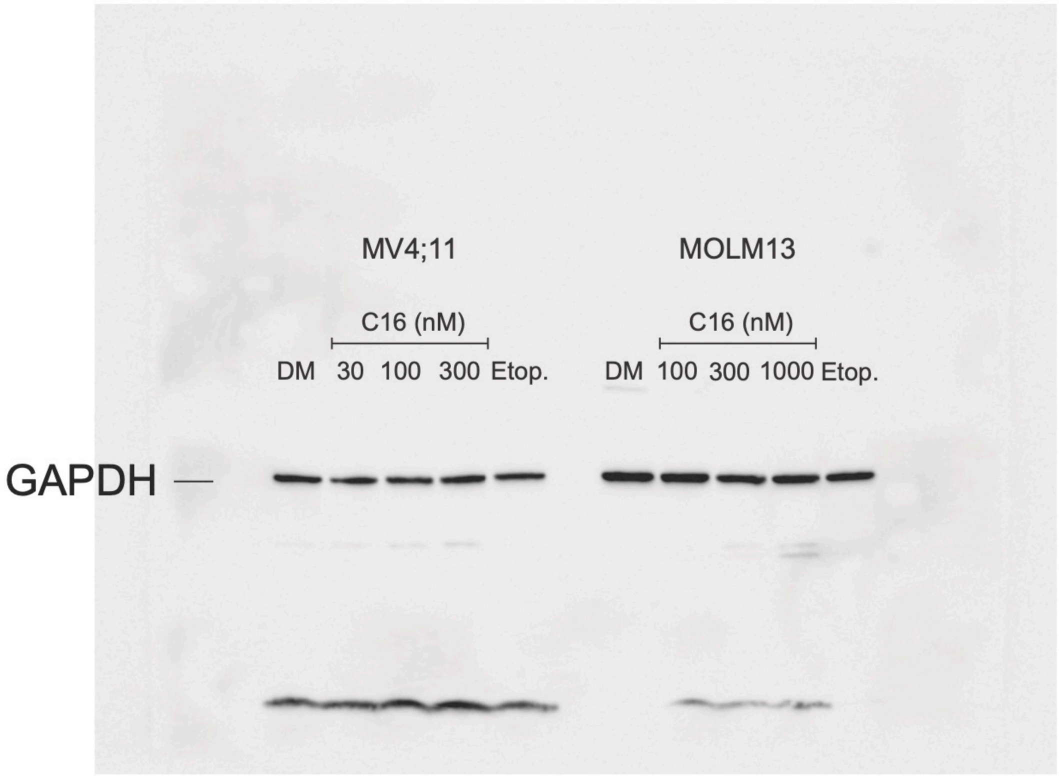

### Figure 6 Source Data 3

**Figure 6—source data 3**

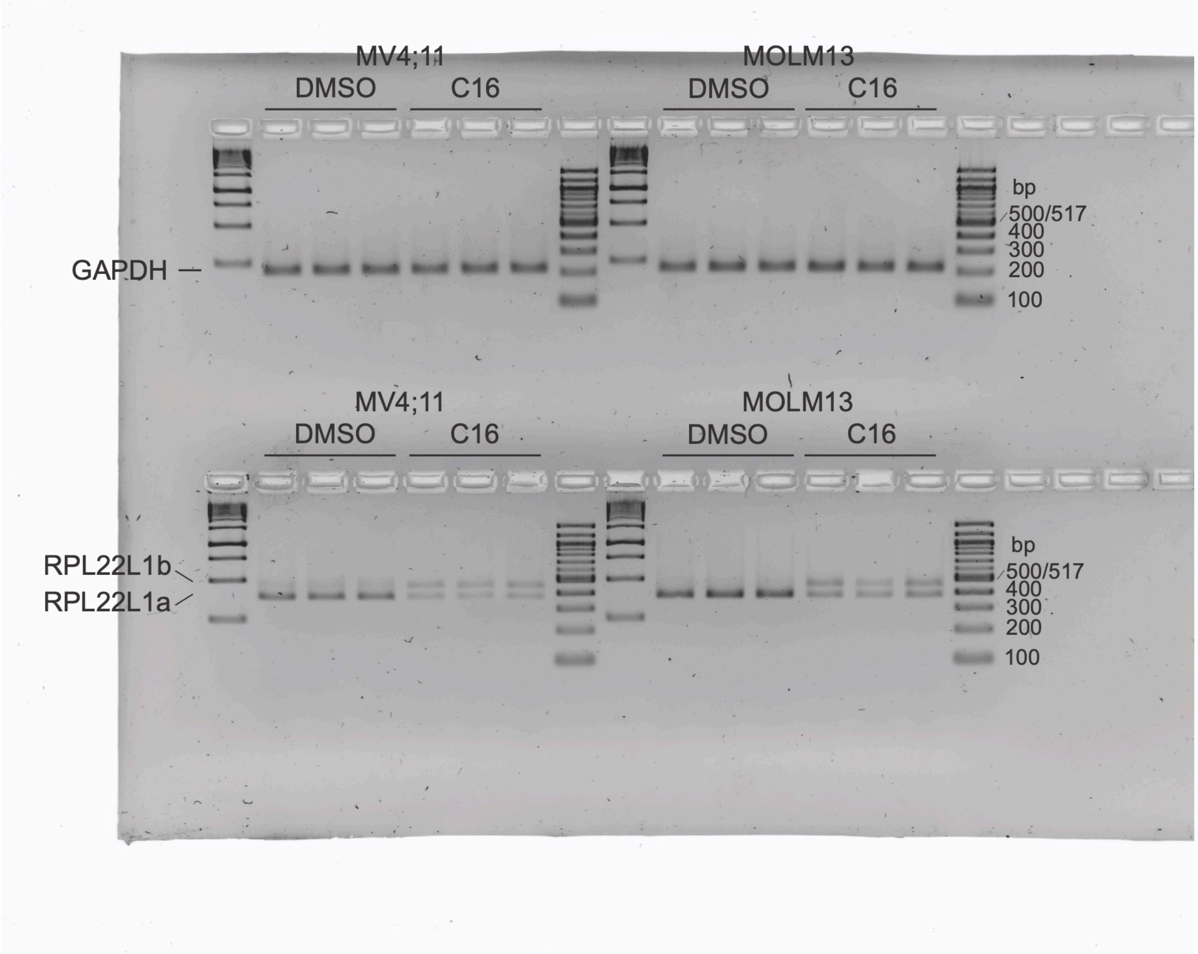


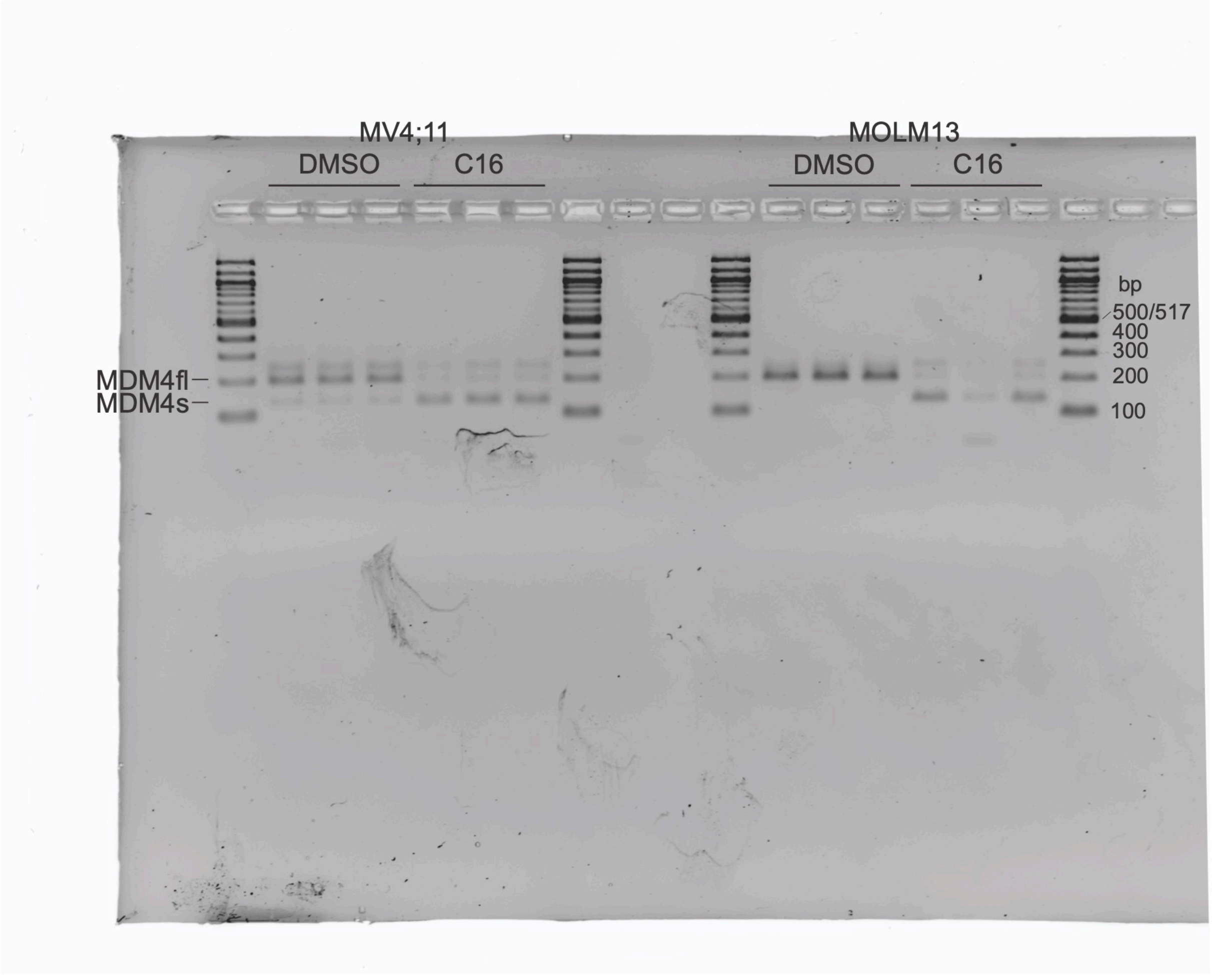

### Figure 6 Source Data 4

##### Figure 6—source data 4

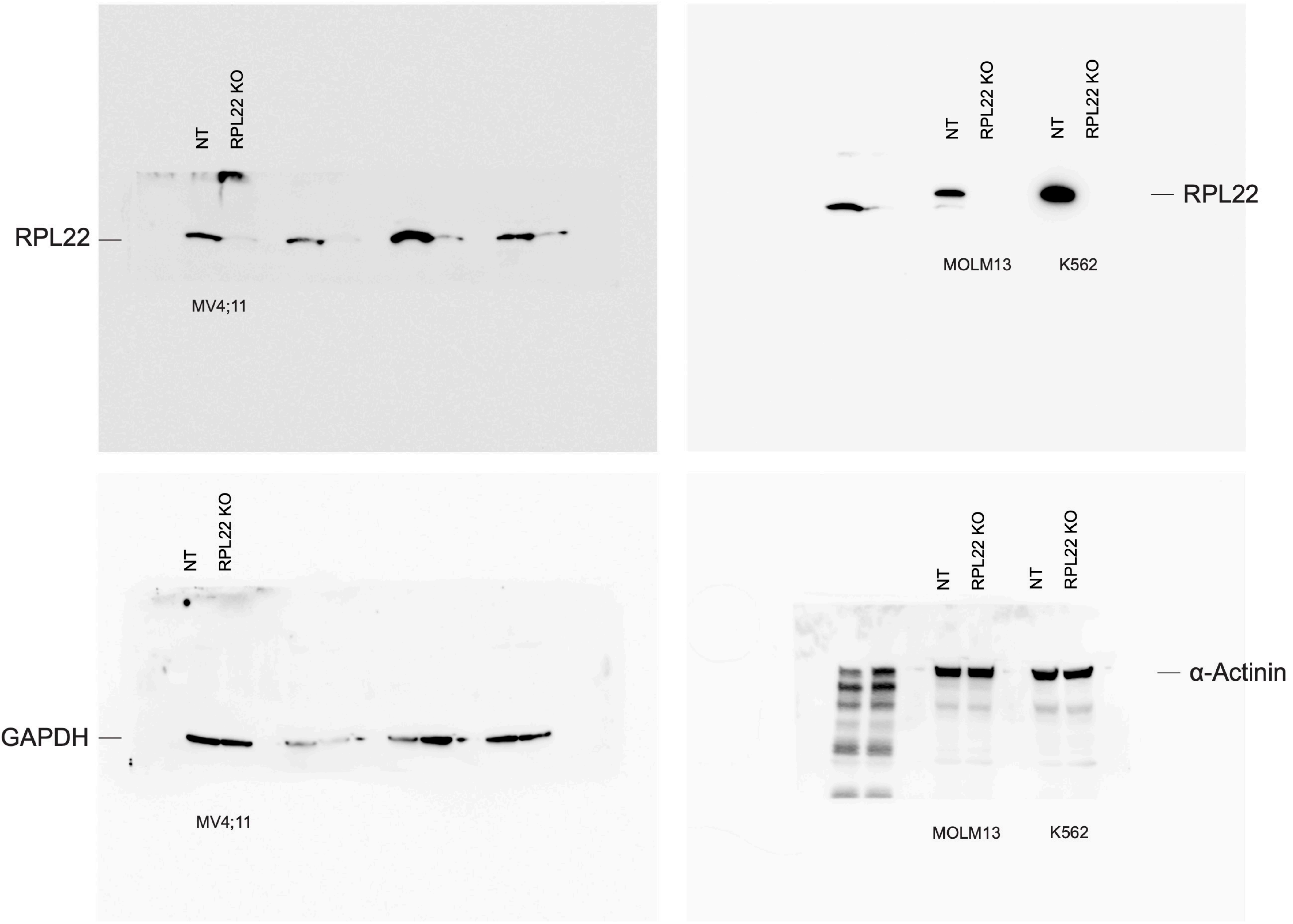

### Figure 6 Source Data 5

#### Figure 6—source data 5

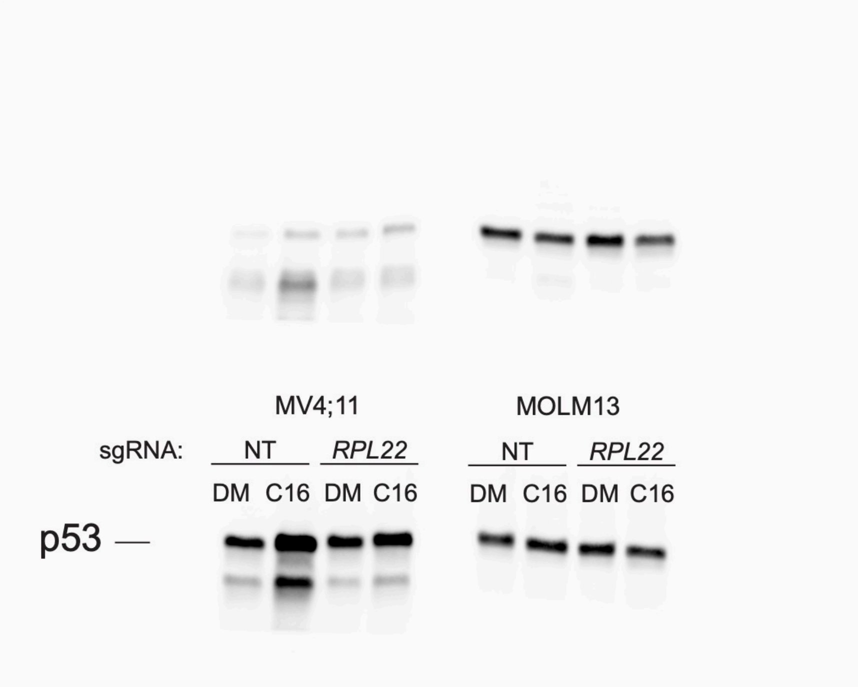


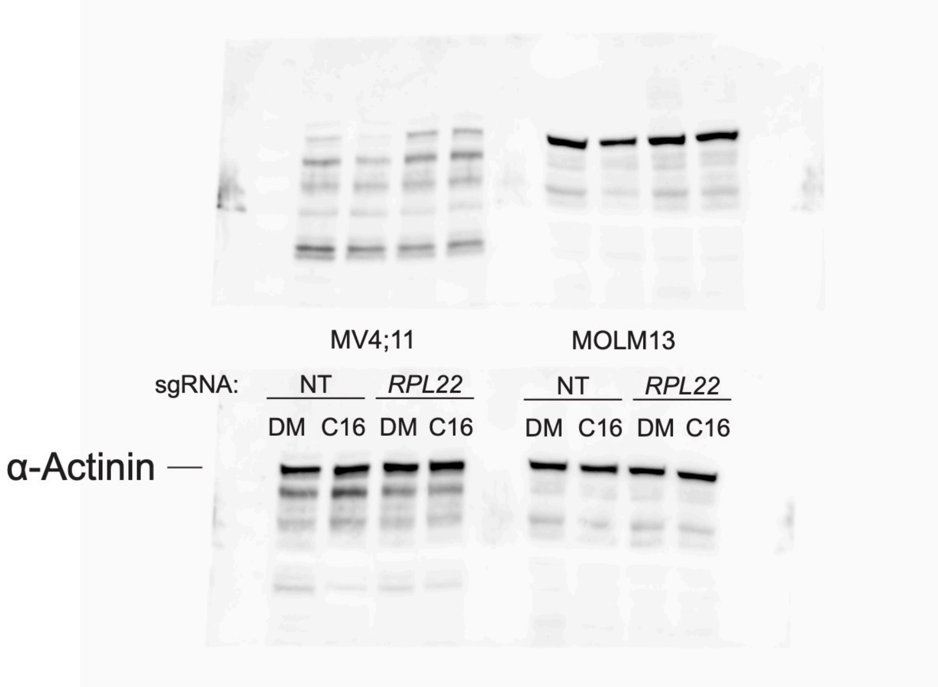

### Figure 6 Source Data 9

**Figure 6—source data 9**

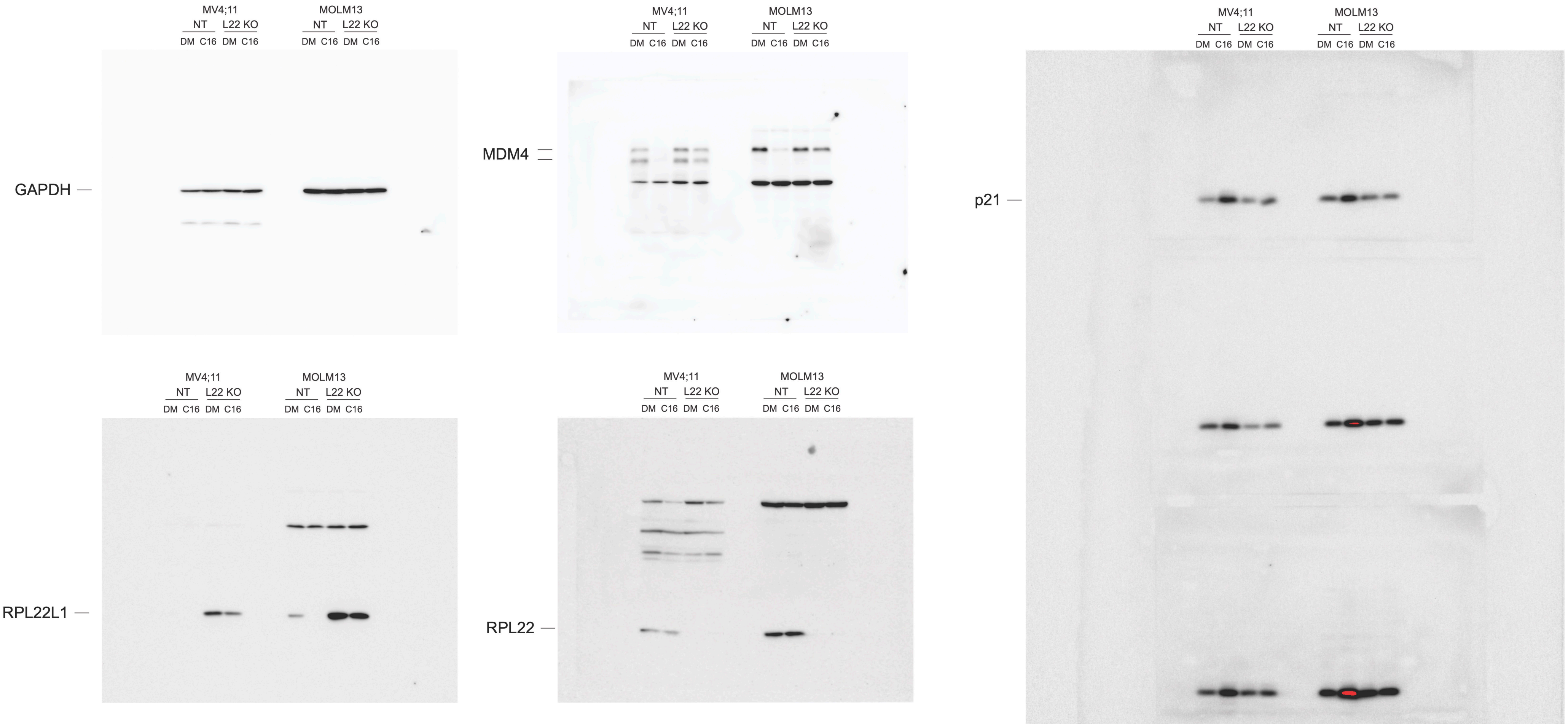
